# sigNATURE maps cohort-specific T-cell states to reproducible programs of ICI response

**DOI:** 10.64898/2026.04.14.718532

**Authors:** Sanika Kamath, Soyeon Kim, Xueke Jin, Jing Hong Wang, Hyun Jung Park

## Abstract

Immune checkpoint inhibitors (ICIs) can induce durable responses across cancers, yet T-cell biomarkers of response remain difficult to reproduce across single-cell RNA-seq studies. A major reason is that T-cell states are typically defined de novo within each cohort, making reported marker genes sensitive to cohort composition and analytic choices rather than stable cellular programs. Here we present sigNATURE (signature Normalization and Atlas-based T-cell Understanding for Reproducibility and Evaluation), a reference-guided framework that maps query cells onto large CD4+ and CD8+ T-cell atlases, evaluates published ICI-response markers in an atlas-aligned coordinate system, and quantifies the atlas support of mapped cells through a cell-level identifiability score. We applied sigNATURE to two independent ICI scRNA-seq cohorts comprising 36 non-small cell lung cancer patients and 15 skin cancer patients (11 basal cell carcinoma and 4 squamous cell carcinoma). Across cohorts, sigNATURE-derived features more robustly resolved response-associated T-cell structure than cohort-derived state definitions, yielding clearer unsupervised separation of responders and non-responders, enabling integrated analysis of independent studies in a shared atlas-aligned space, and improving mean response-prediction AUC from 0.469 to 0.746. Using identifiability score, we further identify terminally differentiated effector CD8+ T cells and regulatory CD4+ T cells as prominent response-associated states across studies, prioritizing published markers in terms of robust, atlas-resolvable cell states. Using this framework. Together, these results establish sigNATURE as a framework for improving the reproducibility, cross-cohort comparability, and mechanistic interpretability of single-cell ICI biomarkers.

## Introduction

Immune checkpoint inhibitors (ICIs) targeting PD-1/PD-L1 and CTLA-4 have transformed cancer therapy by producing durable responses across multiple cancer types. For example, KEYNOTE-024 showed substantial survival benefit with PD-1 blockade in PD-L1-high non-small cell lung cancer (NSCLC) (1), and CheckMate 067 demonstrated long-term benefit of anti-PD-1 alone or in combination with anti-CTLA-4 in metastatic melanoma (2). These therapeutic responses are shaped in part by the composition and functional state of T cells in the tumor microenvironment, including activated, exhausted, and regulatory T-cell populations (3-6). Accordingly, a major goal in cancer immunotherapy has been to identify T-cell states and gene programs associated with clinical response. Several influential single-cell studies have proposed such response-associated programs. For example, Sade-Feldman et al. associated a T-cell subset signature marked by TCF7, IL7R, GPR183, and MGAT4A with response to anti-PD-1- and anti-CTLA-4-based therapies in metastatic melanoma (7), whereas Liu et al. reported that response to anti-PD-1 therapy in NSCLC is linked to expansion of precursor-exhausted CD8+ T cells characterized by high GZMK and relatively low expression of coinhibitory genes such as PDCD1, CTLA4, LAG3, and TIGIT (8). Together, these studies suggest that T-cell response programs may capture biologically meaningful determinants of ICI benefit, but they also raise a central question: do the reported programs reflect conserved features of response across cohorts, or are they substantially shaped by cohort-specific definitions of T-cell state?

Answering this question has been challenging because published single-cell ICI biomarkers often show limited reproducibility across cohorts. Many of these studies are based on modest sample sizes and differ substantially in tumor type, treatment regimen, biopsy timing, and sampling context (9-15). For instance, Sade-Feldman et al. analyzed 32 metastatic melanoma patients across multiple ICI regimens and sampling time points, whereas Liu et al. studied 36 NSCLC patients treated with PD-1-based therapy using paired pre- and post-treatment biopsies. However, heterogeneity in study design is only part of the problem. A more fundamental methodological challenge is that most response-associated signatures are derived from de novo clustering followed by cluster-wise differential expression within each individual cohort. Because cluster boundaries are sensitive to cohort composition, cell-state abundance, and analysis workflow, similar T-cell programs may be partitioned differently across studies (16-18). As a result, reported “response markers” may reflect cohort-specific state definitions as much as conserved, mechanistically meaningful T-cell programs, limiting their transferability across datasets. This makes it difficult to determine whether differences across studies reflect true biological divergence or study-specific partitioning of a shared T-cell state landscape. Moreover, even when query cells are mapped to a reference, most existing analyses provide little indication of how confidently those assignments are supported by the reference landscape, making it difficult to distinguish robust atlas-resolved states from ambiguous mappings near state boundaries (19-22).

To address these challenges, we developed sigNATURE (signature Normalization and Atlas-based T-cell Understanding for Reproducibility and Evaluation), a reference-guided framework for reproducible cross-cohort analysis of ICI-associated T-cell programs. sigNATURE maps cells from published ICI cohorts onto large CD4+ and CD8+ T-cell atlases and benchmarks published response-associated signatures in a shared atlas-aligned coordinate system, enabling direct comparison of T-cell programs across studies. In addition, sigNATURE introduces a cell-level identifiability measure that quantifies how consistently each mapped query cell is supported by its local atlas neighborhood, providing a confidence-aware way to distinguish robust atlas-resolved states from ambiguous assignments. This framework is motivated by the idea that reference-guided mapping can stabilize cellular annotation across studies, while local neighborhood support can reveal whether mapped states are well grounded in the atlas rather than imposed by hard cohort-specific cluster boundaries. Although prior work has shown that bulk transcriptomic predictors can stratify ICI response (23-26), and that reference mapping or graph-based approaches can improve cellular annotation and cross-study comparability (19) (27, 28), these methods do not provide a framework for systematically evaluating published ICI-response signatures in a shared reference-aligned space while simultaneously quantifying the reliability of mapped cellular states. Using two independent ICI single-cell cohorts, we show that sigNATURE improves the reproducibility and cross-cohort comparability of response-associated T-cell features, yields clearer unsupervised separation of responders and non-responders, and improves response discrimination relative to cohort-derived state features. By coupling atlas alignment with identifiability-aware evaluation, sigNATURE provides a principled strategy to distinguish conserved response-associated T-cell programs from cohort-specific state definitions, thereby improving the reproducibility, mechanistic interpretability, and transferability of single-cell ICI biomarkers.

## Results

### Limited reproducibility of published ICI response cell type biomarkers in T cell atlases

To assess the reproducibility of published ICI response T cell markers, we leveraged TCellMap (29), a recently developed T cell reference atlas comprising 308,043 T cells collected from 486 samples of 324 human tissues across 16 cancer types. This atlas provides a comprehensive transcriptional framework for identifying conserved activation and differentiation trajectories of T cells across diverse immune contexts. For focused downstream analysis, we first separated 14 CD8+ T cell states (**Fig. 1A**). Using the CD8+ T cell reference map, we evaluated the expression patterns of ICI-associated markers reported by Sade-Feldman et al. (**Fig. 1B**) and Liu et al. (**Fig. 1C–D**). Although these markers originated from distinct studies and cancer types, we expected them to be localized to similar regions within the reference map, reflecting shared mechanisms of ICI response. Contrary to this expectation, these markers map to different clusters of the CD8+ T cell reference atlas, showing different average expression levels (**Fig. 1E**). Specifically, the Sade-Feldman marker set showed substantial expression (average module score ≈ 0.5) in only 2 of the 14 CD8+ T cell clusters—naïve (Tn) and naïve TCF7+ (Tn_TCF7). In contrast, Liu et al.’s high-expression marker GZMK was broadly expressed across 7 clusters, including transitional effector (t-Teff), central memory (Tcm), proliferative exhausted (p-Tex), senescent (Tsen), and effector states (Teff_CD244□, Teff_SEMA4A□). On the other hand, Liu et al.’s coinhibitory markers such as PDCD1 (PD-1), CTLA4, LAG3, and TIGIT were generally low across the atlas but remained above background in exhausted (Tex) and interferon-response (Tisg) clusters. Together, these observations indicate that ICI-associated markers from different studies map to non-overlapping regions of the CD8+ T cell reference atlas.

**Figure 1.**
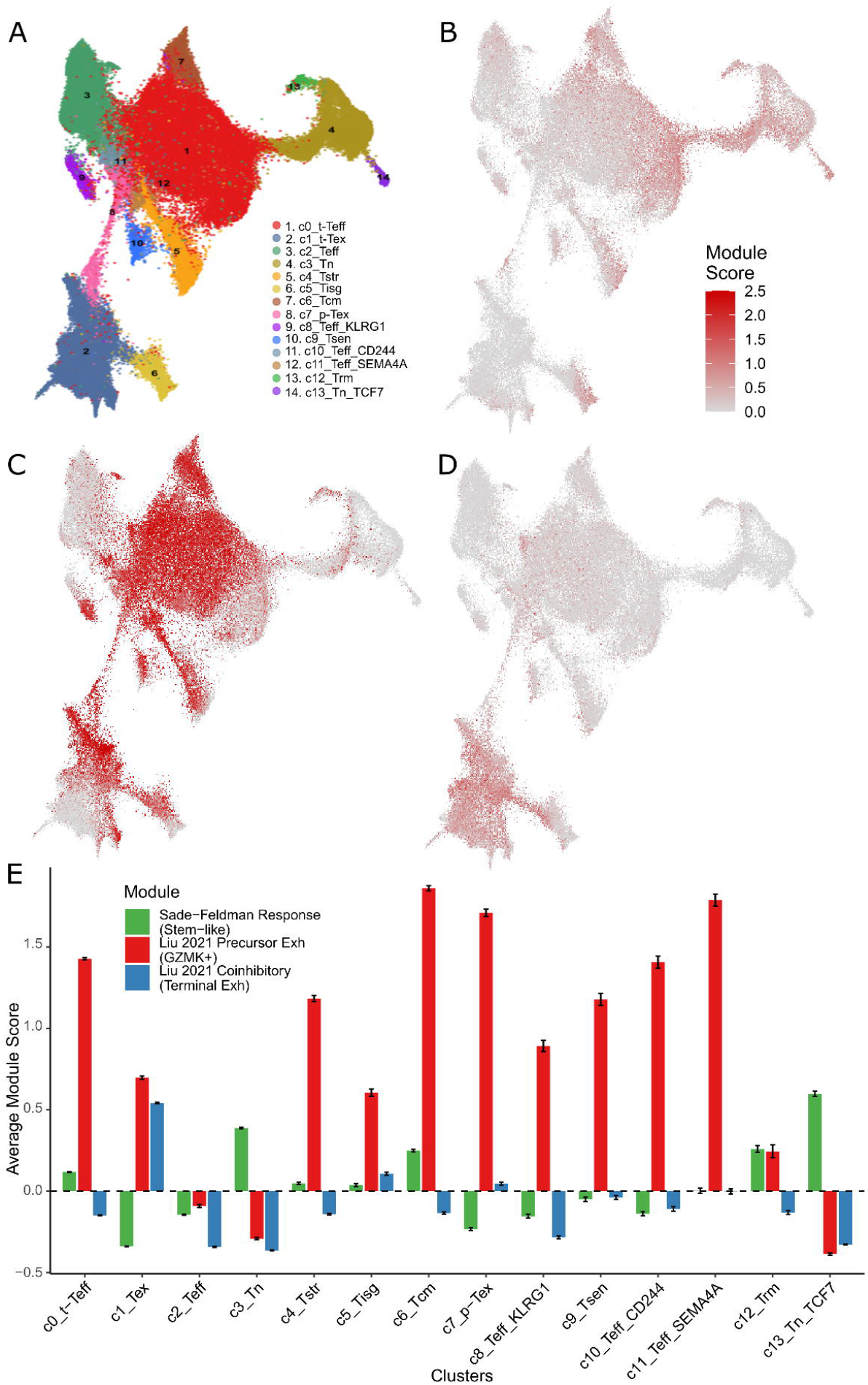
(A) sigNATURE CD8+ cells with cluster definition. (B) Sade-Feldman’s, (C) Liu et al.’s GZMK, and (D) Liu et al.’s coinhibitory markers on sigNATURE CD8+ reference atlas. (E) Average expression value of Sade-Feldman’s (green), Liu’s GZMK (red), and Liu’s coinhibitory (blue) markers across sigNATURE CD8+ T cell clusters.

To further dissect heterogeneity within CD8+ T cell subtype signatures, we examined a curated set of genes implicated in T cell activation, effector function, and immune regulation (**Table 1**). Immediate-early activation markers (FOS, FOSB, CD69) and cytotoxic effectors (GZMA, GZMK) displayed highly variable expression across CD8+ T cell clusters (**S. Fig. 1 A–E**). Similarly, chemokines (CCL4, CCL4L2, CXCL13) and the receptor CXCR6, which mediate tissue trafficking and spatial organization within the tumor microenvironment, also exhibited highly heterogeneous expression patterns (**S. Fig. 1 F-I**). Transcriptional regulators (EOMES) and metabolic/stress modulators (TXNIP, FKBP5) also showed heterogeneous expression (**S. Fig. 1 J-L**). While some markers (e.g., CCL4 and GZMA, CXCR6 and CXCL13) roughly align with each other, genes exhibit overall cluster-specific expression patterns. This variability underscores the complexity of defining robust biomarkers for ICI response and highlights the need for sophisticated approaches that capture functional diversity rather than relying on single-gene or cohort-specific signatures.

**Table 1.**
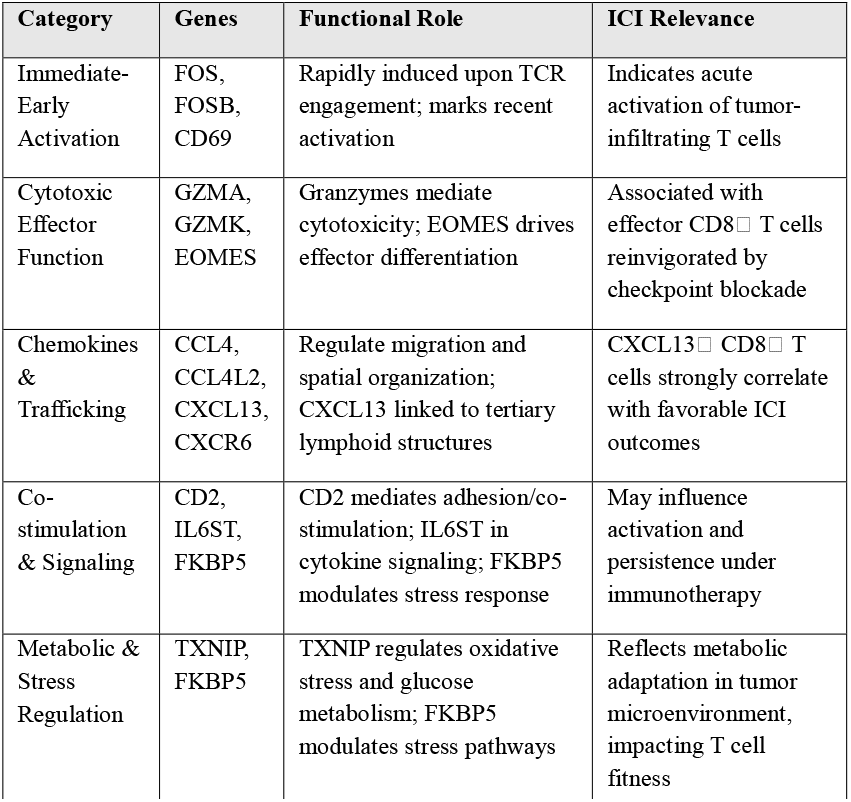
Functional category of genes with ICI relevance.

### Benchmarking ICI-Associated CD8 T Cell Signature Against T Cell Atlas Using sigNATURE Label-transfer

To identify ICI gene markers with functional implication and thus reproducibility, we developed sigNATURE, a systematic framework that utilizes high-resolution CD4+ and CD8+ T cell reference atlases (**Fig. 2A, see Methods**). On these atlases, sigNATURE applies normalization techniques to harmonize heterogeneous datasets and then performs label transfer followed by neighborhood construction to ensure accurate cell identity mapping. To demonstrate this workflow, we applied sigNATURE to the scRNA-Seq dataset from Liu et al (**S. Fig. 2A**). Before performing label transfer, quality control analysis revealed that the Liu et al. cohort showed a systematic difference in cell-cycle activity relative to the reference atlas (**S. Fig. 2A**), with cell-cycle–associated states unevenly distributed across T-cell clusters (**S. Fig. 2B**).

**Figure 2.**
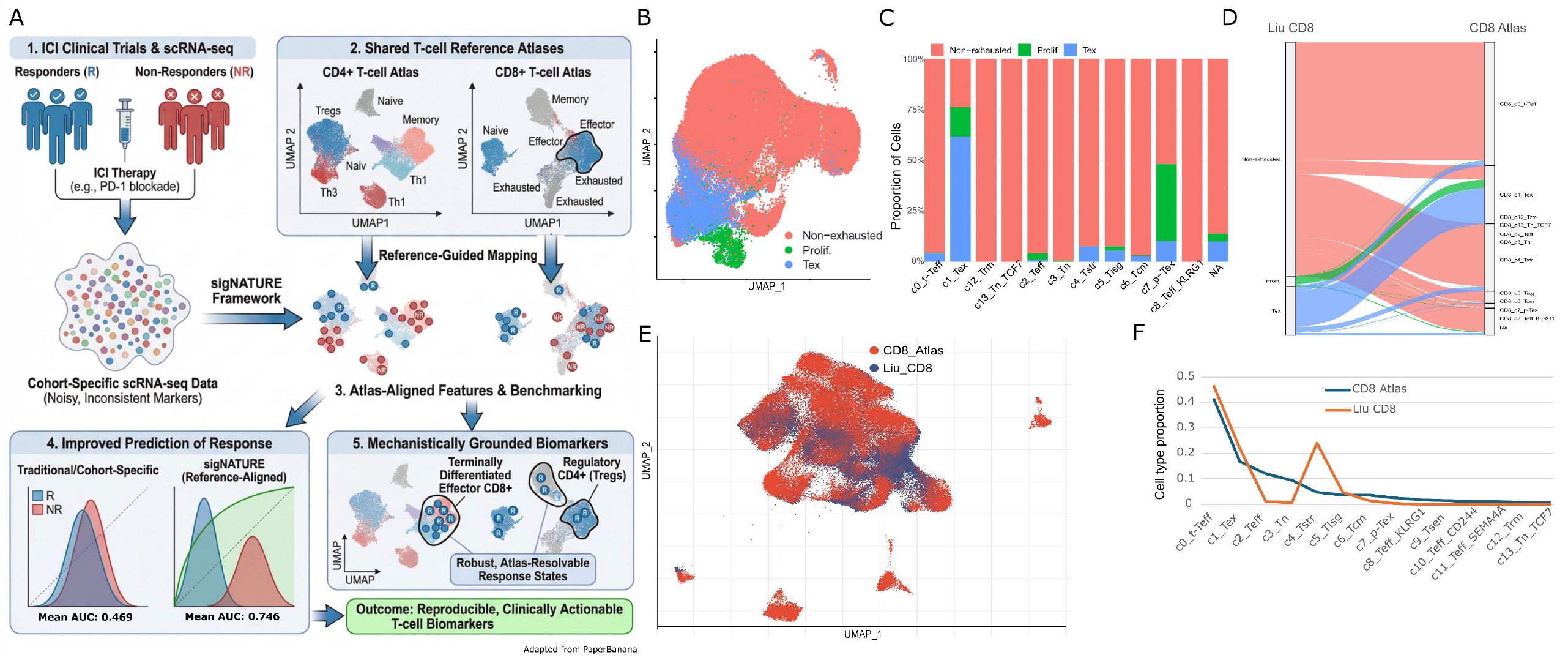
(A) Schematic illustration of sigNATURE that takes single-cell RNA-Seq data of clinical trials to identify reproducible and clinically actionable T-cell biomarkers. (B) UMAP of Liu et al.’s CD8+ T cell clusters after cell-cycle normalization (C) Proportion of T cell types that were defined by Liu et al. on the sigNATURE CD8+ single-cell atlas. (D) Alluvial plot depicting the transfer of Liu et al’s cell clusters to the atlas. (E) Liu et al’s data mapped on the atlas. (F) Cell type proportions in the CD8+ atlas (blue) and Liu et al’s data after mapping (orange).

After regression out the cell-cycle status discrepancy (**S. Fig. 2C, Fig. 2B**), sigNATURE successfully performed label transfer (**Fig. 2C**), and several patterns supported that the mapping was biologically coherent. First, Liu et al.’s non-exhausted cells distributed across several reference states—including transitional effector, effector, stress-response, IFN-response, and central-memory–like states. Also, these reference states are mostly from the non-exhausted cells—consistent with the expectation that naïve/memory/effector phenotypes retain functionality and are not dominated by the chronic antigen stimulation that drives exhaustion. Second, exhausted (Tex) and progenitor exhausted (p-Tex) cells mapped preferentially to exhaustion-related reference states, with minimal leakage into non-exhausted regions (**Fig. 2D**), supporting that the transfer robustly preserved the major functional axis separating non-exhausted from dysfunctional/exhausted programs. Third, sigNATURE adds interpretability by explicitly flagging cells with low mapping confidence as not assigned (NA) (Methods). Notably, some cells originally labeled as non-exhausted in Liu et al. mapped to NA, which may reflect transitional trajectories where boundaries between functional and dysfunctional programs are inherently continuous rather than discrete (30, 31).

A closer look at the projection also highlights resolution limits in Liu et al.’s cohort-defined labels that motivate an atlas-anchored reinterpretation (**Fig. 2E**). First, Liu et al. clustering fails to resolve functionally distinct exhaustion-related states, most notably distinguishing terminally exhausted (Tex) vs. proliferative exhausted T cells (p-Tex) T cells (**Fig. 2F)**. In the atlas, Tex is defined by an exhausted effector program, whereas p-Tex is characterized by strong cell-cycle features, reflecting a distinct functional state (**Fig. 2C**). This illustrates a general limitation of cohort-based marker definition: even if the markers were together highly expressed in responders vs. non-responders in the cohort data, it doesn’t mean the markers represent a unique or coherent functional component of the response mechanism. This discrepancy can lead to false positives or overlook critical markers associated with rare or functionally distinct populations, ultimately compromising the identifiability of biomarker discovery.

### sigNATURE mapping refines CD8 states and improves ICI-response prediction in an independent anti–PD-1 trial

To validate the value of sigNATURE in an independent cohort, we analyzed publicly available single-cell RNA-seq data from a second ICI clinical trial (anti–PD-1 therapy) reported by Yost et al.(32). This dataset includes 11 patients with advanced basal cell carcinoma (BCC) at an average post-treatment sampling time of 54 days. We selected this cohort for validation because it represents a markedly different tumor context and, importantly, because sigNATURE revealed that its cohort-specific structure is not directly portable to atlas-defined cell-state geometry. In the original cohort embedding, CD8+ T cells appear well separated by dataset-derived structure (**Fig. 3A**), consistent with strong within-cohort clustering. However, when the same cells are mapped onto the sigNATURE CD8 reference, they concentrate into a comparatively restricted region of the atlas (**Fig. 3B**), indicating that cohort-driven separations do not necessarily correspond to broadly conserved, reference-resolvable programs. Despite this compression, sigNATURE mapping remained sensitive to biologically meaningful heterogeneity: while the original cohort analysis yielded four broad CD8 compartments (CD8_mem, CD8_act, CD8_eff, CD8_ex), reference mapping followed by sigNATURE-guided refinement resolved nine CD8 neighborhoods of substantial size (**Fig. 3D**). This increase in resolution supports our central premise that atlas mapping can disentangle cohort-specific embeddings into more reproducible, mechanistically interpretable cell states. Finally, we assessed predictive utility in this external dataset using the same pair-based cross-validation framework applied in our primary analysis, which holds out one responder–non-responder pair per fold to provide balanced test sets under small sample sizes.

**Figure 3.**
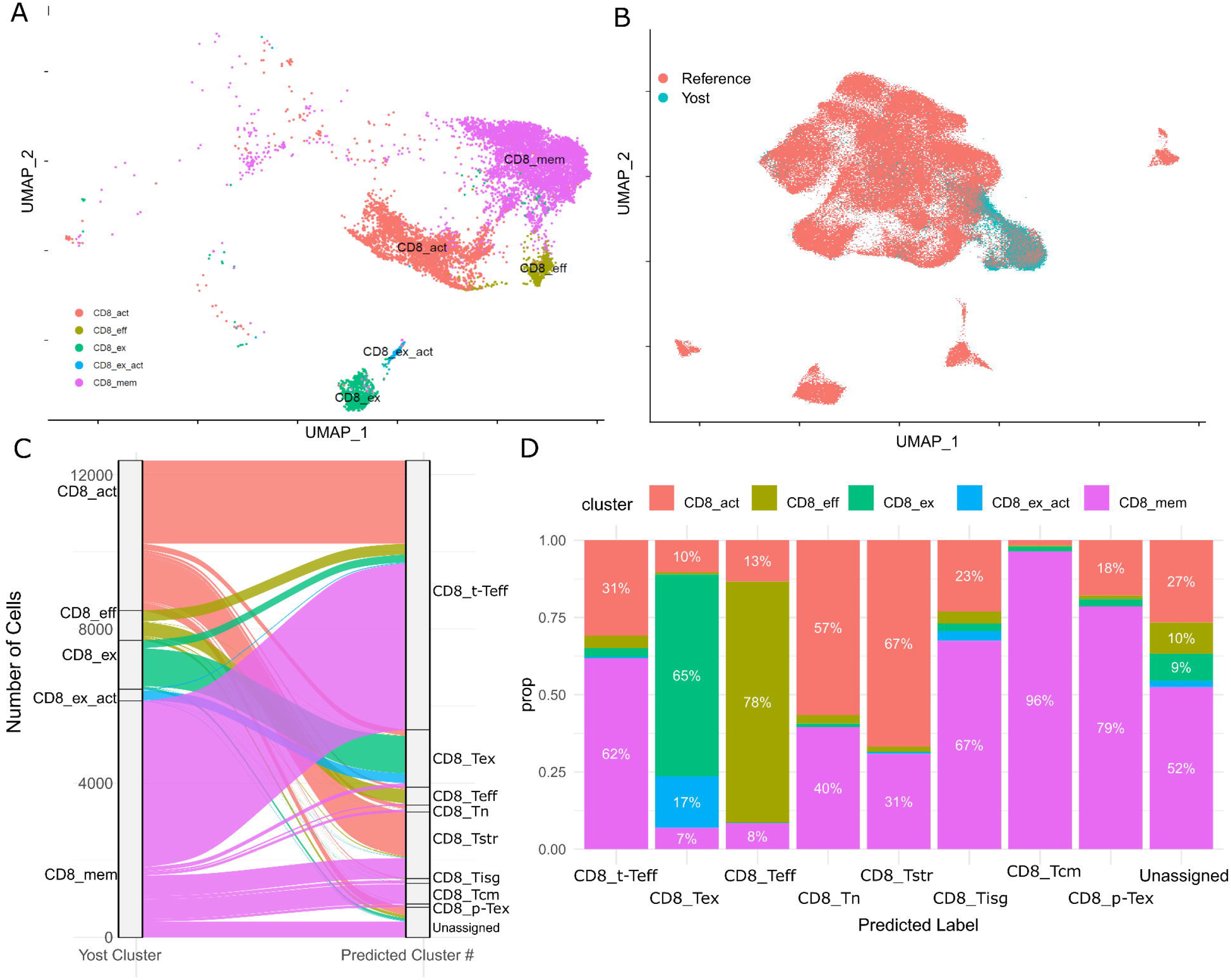
**(A)** UMAP of Yost et al.’s CD8+ T cell clusters as defined in the original work. (B) Proportion of T cell types that were defined by Yost et al. on the sigNATURE CD8+ atlas. (C) Alluvial plot depicting the transfer of Liu et al’s cell clusters to sigNATURE CD8+ atlas.

### sigNATURE reveals reproducible and reference-aligned ICI response programs across cohorts

To determine whether atlas-aligned cell-state features improve the reproducibility of immune checkpoint inhibitor (ICI) response signals, we first analyzed the Liu et al. cohort, in which prior work showed that response-associated clonal expansion is concentrated in exhausted CD8+ T-cell states. Because the sample size is limited (n = 13), we treated supervised prediction in this cohort as exploratory and used it primarily to evaluate whether sigNATURE captures structured response-associated variation. After transferring atlas labels, we quantified sigNATURE-derived CD8 T-cell state proportions and compared their predictive performance with that of the original Liu et al. CD8 state definitions (Methods). The model based on the original cohort-derived states showed minimal discriminatory ability (AUC = 0.472), whereas the sigNATURE-derived model yielded improved discrimination (AUC = 0.528, **Fig. 4A**). Although this gain was modest in absolute magnitude, it was accompanied by a marked difference in how predictive information was distributed across the feature space. To assess whether this gain arose from a small number of dominant states or from broader organization of the feature space, we progressively increased the fraction of sigNATURE features included in the model. Discrimination increased steadily until nearly the full feature set was retained (**Fig. 4B**), and all 27 principal components carried non-negligible predictive information (**Fig. 4C**). These results indicate that sigNATURE encodes response-associated information as a distributed and structured representation of CD8+ T-cell heterogeneity, rather than as a sparse collection of cohort-specific predictors. We next asked whether this more structured representation would translate into more reproducible response discrimination in an independent dataset. In the Yost et al. validation cohort (n = 11 patients), sigNATURE-derived cell-state proportion features markedly outperformed the corresponding cohort-derived CD8 feature set in discriminating responders from non-responders (AUC = 0.967 versus 0.467, **Fig. 4D**). Despite a similar sample size limitation, the consistent improvement across the cohorts suggests that sigNATURE is not simply reparameterizing immune heterogeneity within a single study, but is instead recovering response-associated cellular programs that are more stable across datasets.

**Figure 4.**
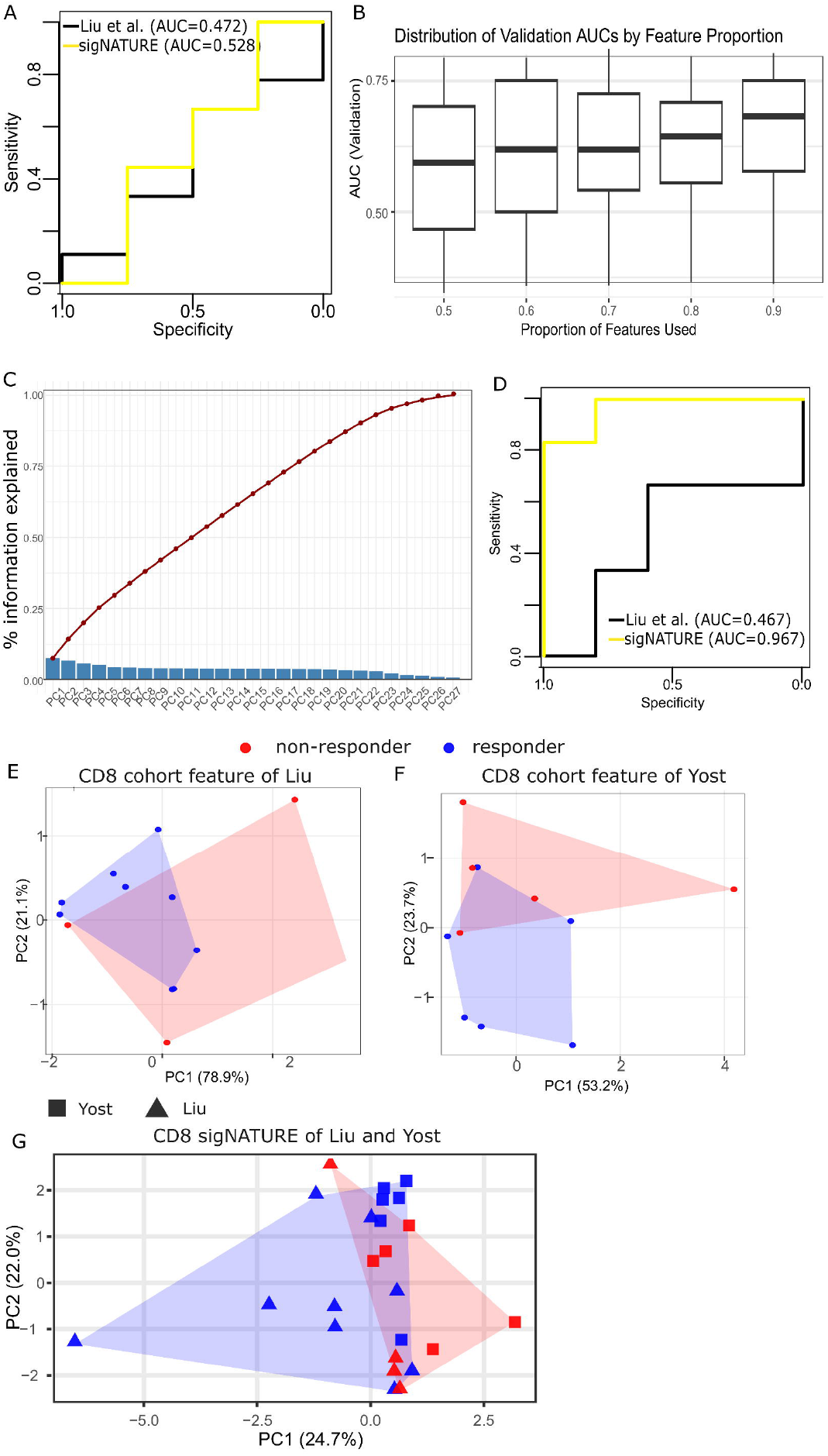
**(A)** Predictive performance in the Liu et al. cohort (n = 13), comparing response discrimination based on cohort-derived CD8+ T-cell state proportions versus sigNATURE-derived CD8+ T-cell state proportions. Because of the limited sample size, this analysis was treated as exploratory. **(B)** AUC of the response-prediction model in the Liu et al. cohort as progressively larger fractions of sigNATURE features were included, showing that predictive information is distributed broadly across the feature space rather than concentrated in a small subset of dominant features. **(C)** Contribution of each principal component (PC) of the sigNATURE feature space to response prediction in the Liu et al. cohort, showing that predictive information is distributed across the full representation. **(D)** Predictive performance in the independent Yost et al. validation cohort (n = 11), comparing response discrimination based on cohort-derived CD8+ T-cell state proportions versus sigNATURE-derived CD8+ T-cell state proportions. **(E, F)** Dimensionality reduction of cohort-derived T-cell features in the Liu and Yost cohorts, respectively, showing limited separation of responders and non-responders. Convex hulls indicate the distribution of samples by response group. **(G)** Dimensionality reduction using sigNATURE-derived features, showing clearer separation of responders and non-responders and demonstrating that atlas-aligned features better resolve response-associated immune structure than cohort-derived state definitions.

Further, we sought complementary evidence from unsupervised analyses. In separate analyses of the Liu and Yost cohorts, dimensionality reduction of cohort-derived CD8 or CD4 T-cell features showed substantial intermixing of responders and non-responders, with overlapping convex hulls in the projected space (**Fig. 4E, 4F, S. Fig. 3A, 3B**). By contrast, sigNATURE-derived features showed a stronger tendency for responders and non-responders to occupy different regions of the embedding, both within cohorts (**S. Fig. 3C**) and in joint analysis across cohorts (**Fig. 4G**). Because sigNATURE projects cells from independent studies into a shared reference-aligned state space, it enabled direct joint analysis of the Liu and Yost cohorts in a common low-dimensional representation. Relative to separate cohort-level embeddings, this combined projection places more samples into the same response-aligned landscape, making regions enriched for responders or non-responders more apparent in the shared space (**Fig. 4G**). Although these patterns should be interpreted cautiously given the modest sample sizes and the limitations of two-dimensional visualization, they are consistent with the supervised analyses in suggesting that sigNATURE organizes response-associated variation in a manner that is more comparable across datasets than cohort-derived state features. Taken together, these results support the conclusion that sigNATURE improves ICI response analysis by transforming cohort-specific immune states into more reproducible, reference-aligned cellular programs.

### sigNATURE Prioritizes ICI-Response T-Cell Markers Using Its Identifiability Measure

To identify ICI-response T-cell markers with clinical utility, we introduce a cell-level identifiability score that quantifies how consistently each mapped query cell is supported by its local atlas neighborhood. After projecting a query cell into the atlas embedding, we retrieve its *k* nearest atlas neighbors and compute identifiability as the proportion of those neighbors that fall within the selected top atlas clusters in that neighborhood. This score increases when the cell’s nearest neighbors are dominated by a coherent atlas cluster signal (high identifiability) and decreases when neighbors are distributed across multiple clusters (low identifiability) (**Fig. 5A**). Based on this measure, sigNATURE revealed distinct identifiability of CD8 T cell states in the Liu et al’s data. For example, Tstr and Tisg are marked by distinctive, high-amplitude programs (heat-shock/stress genes for Tstr; interferon-stimulated genes for Tisg), which makes them readily separable from neighboring states (33). Likewise, Tex is defined by a coordinated exhaustion module (e.g., PDCD1/TIGIT/LAG3/HAVCR2 with TOX) (33). In contrast, Tcm and p-Tex do not form sharply defined atlas states, but instead occupy transitional regions that bridge effector and exhausted neighborhoods—consistent with p-Tex acting as a continuum state in pan-cancer atlases. As a result, their boundaries are diffuse rather than discrete, leading to lower identifiability despite apparent enrichment (33). Traditionally, biomarkers were identified mostly based on enrichment magnitude in responders vs. non-responders. Under that criterion, the top CD8+ candidates would be Teff_KLRG1, Trm, Tn_TCF7, Tn, and terminal effector (t-Teff) (**Fig. 5B**). However, by imposing the constraint of atlas-based identifiability, sigNATURE isolates t-Teff as the sole top-ranked candidate with robust identifiability, while filtering out others due to ambiguity or overlap.

**Figure 5.**
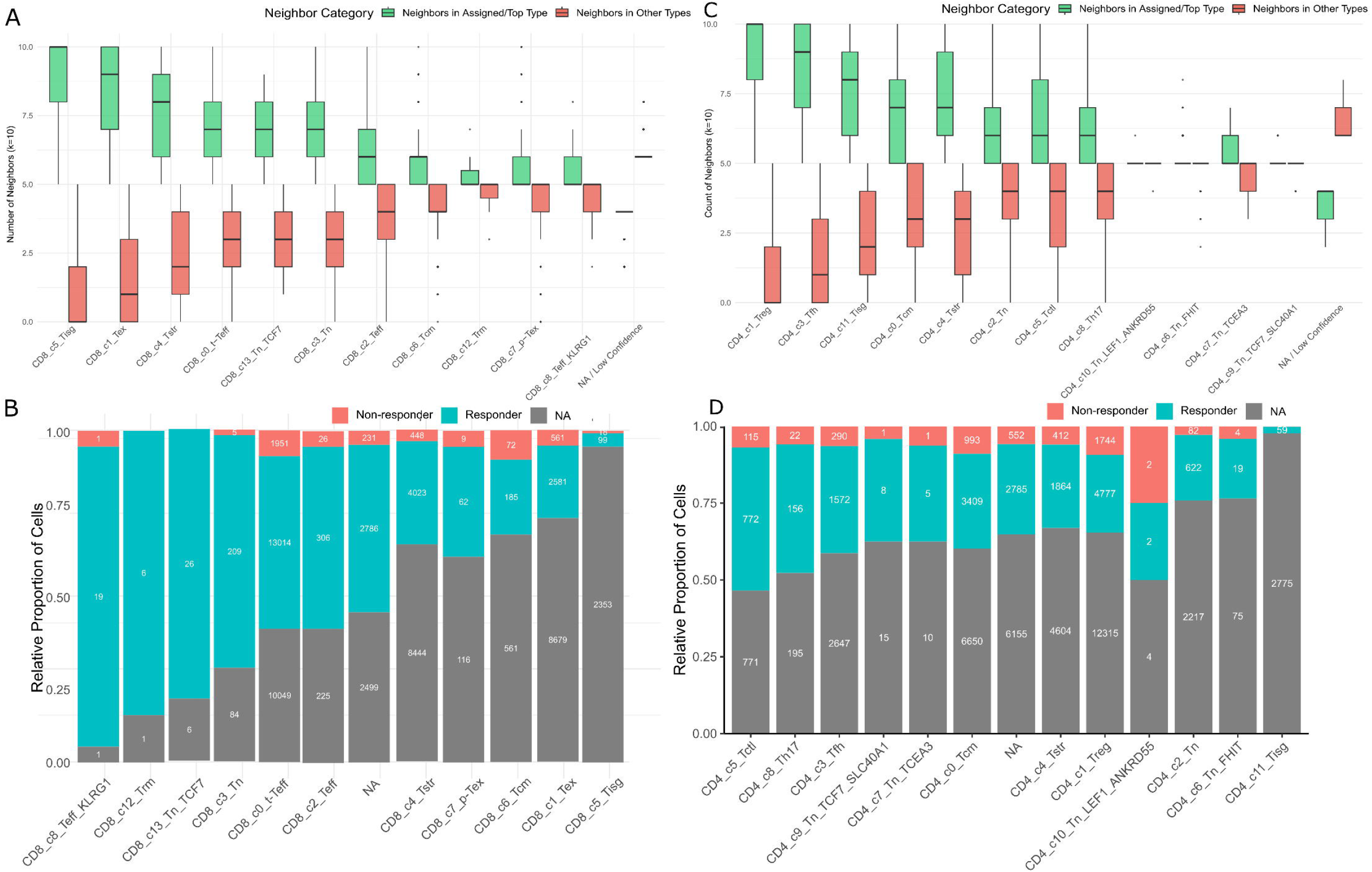
Evaluation of mapping stability and association with response. (A, C) Box plots quantifying mapping stability for (A) CD8+ and (C) CD4+ T cell states. Stability is measured by the number of nearest atlas neighbors (k=10) that share the cell’s assigned cluster (green) versus those belonging to different clusters (red). (B, D) Relative proportion of (B) CD8+ and (D) CD4+ sigNATURE subtypes, including the ‘Not Assigned’ (NA) category and stratified by clinical outcome. Bars represent the proportion of cells from Responders (R) versus Non-responders (NR) within each subtype. The cell-subtypes are ranked by their enrichment in Responders.

Similarly, among Liu et al.’s CD4 T cell states, Treg, Tisg, and Tfh tend to map more confidently (**Fig. 5C**). They are characterized by strong, lineage-defining transcriptional programs that recur across studies. For example, pan-cancer atlas work resolves a CD4 Treg state with canonical IL2RA/FOXP3/CTLA4 expression and a Tfh state marked by ICOS, BCL6, CXCL13, and PDCD1, alongside a coherent IFN-response (Tisg) program (33). Using the same logic as above, Treg shows the strongest enrichment in responders vs. non-responders while also mapping confidently to the reference atlas. Therefore, CD4 Treg emerges as a strong candidate ICI-response subset, supported by both identifiability and responder enrichment.

## Discussions

Immune checkpoint inhibitors (ICIs) have transformed cancer therapy, yet biomarkers that reliably predict response remain elusive. Although numerous single-cell studies have reported ICI-associated T-cell states and proposed marker-gene signatures, these signatures often fail to generalize across cohorts, cancer types, and profiling protocols. A key methodological reason is that most signatures are derived by de novo, cohort-specific clustering followed by cluster-wise differential expression, such that reported “response markers” can reflect dataset-specific state boundaries and sampling context as much as stable, mechanistically meaningful T-cell programs (16-21). To address this challenge, we present sigNATURE, a reference-guided framework that anchors cells to unified CD4+ and CD8+ T-cell atlases, leverages atlas-neighborhood structure, and evaluates both response enrichment and identifiability. By doing so, sigNATURE converts cohort-specific single-cell findings into reference-aligned cellular programs that are more reproducible and biologically interpretable across studies.

By applying sigNATURE to multiple published ICI datasets, we show that many previously reported response-associated markers do not converge on conserved T-cell states when examined in a unified reference framework. Instead, these markers map to heterogeneous and only partially overlapping regions of the CD4+ and CD8+ T-cell landscapes, highlighting a fundamental limitation of cohort-derived biomarker discovery (**Fig. 1**). Together with the many-to-many correspondence between de novo clusters and atlas states (**Fig. 2**), these findings indicate that biomarker instability often arises not only from biological variability, but also from cohort-specific state definitions. In this setting, sigNATURE provides a principled alternative by reinterpreting response-associated signals relative to a shared cellular reference rather than to cohort-restricted clusters alone. Importantly, the identifiability measure strengthens this\ framework by quantifying how consistently mapped query cells are supported by their local atlas neighborhood, thereby helping distinguish robust atlas-resolvable states from more ambiguous assignments near state boundaries.

Our analyses further show that the value of sigNATURE extends beyond improved state annotation or modest gains in supervised prediction. In the Liu cohort, where the limited sample size necessitates caution in interpreting classification performance, sigNATURE nevertheless captured response-associated information as a structured, distributed representation rather than as a sparse set of dominant features (**Fig. 4A–C**). This interpretation was supported by complementary unsupervised analyses: whereas cohort-derived CD4+ and CD8+ T-cell features showed substantial responder/non-responder overlap in the Liu and Yost cohorts, sigNATURE-derived features produced embeddings in which response groups were more spatially organized, both within cohorts and in joint analysis across cohorts (**Fig. 4E–G; Supplementary Fig. 3**).

Although such visual patterns should be interpreted cautiously, they are consistent with atlas alignment organizing response-associated variation in a manner that is less dependent on cohort-specific cell-state definitions. The independent Yost cohort further supports this view. Although both Liu and Yost remain relatively small for definitive predictive modeling, sigNATURE-derived features improved response discrimination in the validation cohort and did so in the same direction as the broader structural patterns observed in unsupervised analyses. Moreover, because sigNATURE projects independent datasets into a shared reference-aligned space, it enables joint cross-cohort analysis of response-associated immune states; in our joint embedding, combining both studies made responder-enriched and non-responder-enriched regions more apparent than in separate cohort-level analyses. Thus, the primary contribution of sigNATURE is not simply higher classification accuracy in a small dataset, but the ability to transform unstable cohort-specific immune states into reproducible, reference-aligned programs that can be compared, interpreted, and integrated across studies. This distinction is important because, in small single-cell clinical datasets, unsupervised structural coherence and cross-cohort consistency may provide more reliable evidence of biological relevance than isolated performance gains alone (34). In this setting, identifiability serves as a useful confidence-aware complement by prioritizing response-associated states that are not only enriched or predictive, but also stably supported by the atlas structure.

Several limitations warrant further consideration. First, although the current CD4+ and CD8+ reference atlases provide broad coverage of T-cell transcriptional programs, sigNATURE will likely improve as larger, more diverse, and more context-rich atlases become available (19-21). This is relevant not only for atlas alignment itself, but also for identifiability, because the precision of local neighborhood support depends on the density and diversity of the reference landscape. Second, our analyses focused primarily on transcriptomic states; integrating chromatin accessibility, T-cell receptor, protein, and spatial information may further refine state definitions and enhance mechanistic interpretation (35-37). Third, while sigNATURE improved both supervised and unsupervised resolution of response-associated structure, the predictive analyses were performed in relatively small cohorts, limiting the precision and stability of cross-validated AUC estimates (34). Future studies in larger and more diverse ICI datasets will be needed to more fully assess predictive generalizability and to establish how atlas-aligned cellular programs relate to clinical response across tumor types and treatment settings.

More broadly, sigNATURE establishes a general paradigm for translating single-cell discoveries into clinically meaningful biomarkers by grounding them in a shared, biologically interpretable reference (19-21, 38). Rather than asking whether a specific gene or cluster predicts response in a single cohort, sigNATURE reframes the question as which conserved cellular programs are reproducibly engaged by therapy across studies and contexts. This shift has important implications for biomarker development, patient stratification, and trial design, because it provides a more stable basis for cross-study comparison, reduces false discoveries driven by technical or cohort-specific effects, and highlights cell states that are more plausibly linked to therapeutic efficacy. The central message is that reproducible and clinically actionable biomarkers are more likely to emerge from reference-aligned cellular programs than from isolated cohort-specific gene signatures, and sigNATURE provides a practical framework for achieving that goal.

## Figure Captions

**S. Fig. 1** Expression UMAP of (A) FOS, (B) FOSB, (C) CD69, (D) GZMA, (E) GZMK, (F) CCL4, (G) CCL4L2, (H) CXCL13, (I) CXCR6, (J) EOMES, (K) TXNIP, (L) FKBP5 on sigNATURE CD8+ reference atlas.

**S. Fig. 2** (A) Cell proportion of Liu et al’s and our CD8 reference data in terms of cell cycle information, whether it is G1, G2M, or S. (B) Cell proportion of Liu et al’s CD8 T cell states in terms of cell cycle information, whether it is G1, G2M, or S. (C) UMAP of Liu et al.’s T cell clusters before cell-cycle normalization

**S. Fig. 3 (A, B)** Low-dimensional projections of cohort-derived CD4+ or CD8+ T-cell features from the Liu et al. and Yost et al. cohorts, showing substantial overlap between responders and non-responders. **(C)** Low-dimensional projection of sigNATURE-derived features showing improved separation of response groups and enabling direct comparison of independent cohorts in a shared atlas-aligned state space.

## Materials and Methods

### Data Acquisition

To evaluate the reproducibility of immune checkpoint blockade (ICB) response markers and characterize the functional states of T-cells, we utilized two pre-existing single-cell RNA sequencing (scRNA-seq) datasets. The query dataset was obtained from Liu et al. (2022) via the Gene Expression Omnibus (GEO; accession number GSE179994). This dataset comprises 59,656 CD8+ and 75,735 CD4+ T-cells isolated from 47 biopsies collected from 36□patients with non-small-cell lung cancer (NSCLC). There were 33 pre-treatment and 14 post-treatment samples with anti-PD1 ICB therapy. Clinical metadata provided by the authors allowed me to stratify these samples into two binary clinical outcomes: Responders (9), Non-Responders (4), and others (unknown, designated as ‘-’).

The Liu et al. cohort had dataset-specific clustering to define the T-cell states. We projected these cells onto a standardized, high-resolution baseline. For this structural framework, a comprehensive T-cell reference atlas from (Chu et al., 2023) was used. This atlas comprises 308,043 T-cells (110,218 CD8+ T-cells and 171,761 CD4+ T-cells) collected from 486 samples from 324 individuals across 16 human cancer types, capturing conserved activation and differentiation trajectories across diverse immune contexts.

### Gene Signature Module Scoring

To evaluate the functional gene signature expression and test whether literature-derived biomarkers map to consistent functional states on a unified reference, we used the AddModuleScore function in Seurat to evaluate the functional gene signature expression on the reference dataset. Prior to scoring, data layers were joined to ensure consistent calculation across batches. Three specific gene modules were defined to characterise T-cell states:

1. Response Genes: *TCF7, IL7R, GPR183, MGAT4A* (Sade-Feldman et al.).
2. Precursor Exhaustion: *GZMK* (Liu et al.).
3. Terminal Exhaustion: *PDCD1, CTLA4, LAG3, TIGIT* (Liu et al.).

To compare the expression patterns across the T-cell states, we visualized the module scores for all three defined gene modules on the combined UMAP embedding using the FeaturePlot function in Seurat. A custom color gradient (Gray to Red) was applied with fixed limits (0 to 2.5) to ensure visual comparability across modules; values exceeding the upper limit were squished. Additionally, mean module scores were calculated for each cluster. These were visualized as bar plots with error bars representing the Standard Error (SE) of the mean.

To further dissect heterogeneity within CD8+ T cell subtype signatures, we examined a curated set of genes implicated in T cell activation, effector function, and immune regulation (**Table 1**).

### Query Data Preprocessing and Cell Cycle Correction

Prior to mapping the query dataset (Liu et al., GSE179994) onto the reference, necessary preprocessing steps were performed to mitigate the effects of cell proliferation on clustering. We calculated cell cycle phase scores on the query dataset based on a defined set of G1/S and G2/M markers as described by (Tirosh et al., 2016). This was performed using the CellCycleScoring function in Seurat, following the standard workflow outlined in the Seurat “Cell-Cycle Scoring and Regression” vignette. The difference between the S and G2M scores was regressed out during the ScaleData step to preserve biological heterogeneity independent of cell cycle status.

### Reference-Based Integration and Projection

The query dataset, Liu et al. (GSE179994), was mapped onto this reference using the Seurat FindTransferAnchors and MapQuery functions. Anchors are essentially pairs of cells between the reference and query datasets that exhibit shared biological states. Transfer anchors were identified in PCA space (dims 1:20), and cell type labels were transferred from the reference to the query.

Simultaneously, the query cells were projected into the reference UMAP geometry (ref.umap) to facilitate direct spatial comparison.

To visualize the reference and query datasets together,we built a composite UMAP embedding by joining the original UMAP coordinates of the reference cells with the projected UMAP coordinates of the query cells. This approach allowed us to show the query cells within the stable structure of the reference data without recalculating the manifold. For cluster annotations, we combined the original reference cell types with the predicted labels for the query cells.

### k-NN Consensus Classification and Evaluation of Mapping Confidence

This method introduces a custom “majority vote” classification step using k-NN which is distinct from the standard Seurat prediction method.

#### Feature Alignment and Preprocessing

To ensure exact feature space compatibility between the reference and query datasets, a custom imputation function was developed. Genes present in the reference atlas but absent in the query dataset (Liu et al., GSE179994) were added to the query matrix with zero-expression values. Both the reference CD8+ T-cell atlas and the query dataset underwent standard normalization and scaling. PCA and UMAP models were constructed for the reference object to serve as the structural framework for integration.

### Reference projection based on k-NN Classification

To refine cell type assignment, a custom k-Nearest Neighbour (k-NN) consensus classifier was implemented using the FNN package. For every query cell projected into the reference PCA space, the 10 nearest neighbors from the reference dataset were identified. A majority vote rule was applied to assign labels: a specific cell type was assigned only if at least 5 of the 10 neighbours shared the same annotation. Cells failing to meet this consensus threshold were designated as unclassified (NA).

### Visualization of Cluster Correspondence

The relationship between the original clusters defined in Liu et al. and the k-NN consensus labels assigned by the sigNATURE framework was visualised using an alluvial plot through the ggalluvial package (Brunson, 2020). This plot illustrates the flow of cells from their source identity to their mapped identity. Additionally, the distribution of predicted labels, including unclassified cells, was quantified and visualized using bar plots (ggplot2).

### Evaluation of Mapping Confidence

To go beyond responder/non-responder enrichment alone, sigNATURE introduces an Identifiability criterion that directly tests whether a projected cell (and thus the signature-high region it occupies) corresponds to a coherent atlas-defined T-cell state: for each projected ICI cell **q**, Identifiability is computed as the fraction of its k=10 nearest atlas neighbours whose atlas cell-type label matches the label transferred to the query cell.

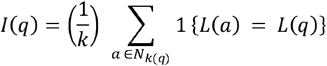

where **{·}** is the indicator function (equals 1 if the condition is true and 0 otherwise) and refers to the cell identity as defined in the embedding. High Identifiability indicates that a published signature localizes to a well-defined atlas state, whereas low identifiability flags signatures that are diffuse across states or sensitive to cohort-specific structure, providing a mechanistic explanation for why some published markers fail to reproduce across studies even when they appear enriched in responders in their original cohort.

To visually evaluate the reliability of these k-NN classifications across various T-cell states, the resulting distributions were depicted through comparative boxplots. For each query cell, the count of nearest neighbors corresponding to the assigned label was compared to the number of non-matching neighbors. These distributions were arranged by the median difference between matching and non-matching neighbors, offering a clear visual assessment of mapping accuracy for each cluster.

### Predictive Modeling

#### Feature Aggregation

To evaluate the clinical utility of the derived cell annotations, single-cell data were aggregated to the patient sample level. For each sample, the cellular composition was quantified by calculating the frequency of each cell cluster relative to the total CD8+ T-cell count in that sample. A binary clinical outcome was defined based on the patient’s response status (Responder = 1, Non-Responder = 0).

Multivariate logistic regression models were constructed to predict clinical response utilizing the aggregated cellular compositions. Two distinct feature sets were evaluated:

#### Baseline Model (Liu Clusters)

Predictors consisted of the proportions of the original broad clusters (*Non-exhausted, Proliferating, Tex*).

### Granular model

Predictors consisted of combinatorial cell states derived by intersecting the original cohort annotations with the high-resolution reference atlas mapping. This approach explicitly captures the hidden heterogeneity within the original broad annotations (for example, specific sub-population of original Liu “Tex” cells that mapped to the reference “Trm” state). Because this intersection maps the original broad clusters across the high-resolution atlas, it expanded the feature space to 27 distinct granular cell states.

For each patient sample, every unique intersecting state was included as a continuous variable representing its fractional proportion of the total CD8+ T-cell count. Because these aggregate proportions sum to 1.0 within each sample, they introduce strict multicollinearity. During model fitting, the glm algorithm inherently accounts for this compositional constraint, and the high feature-to-sample ratio, by automatically dropping collinear covariates to ensure a mathematically valid matrix inversion.

Model performance was assessed using receiver operating characteristic (ROC) curve analysis. The area under the curve (AUC) was calculated for each model to quantify its discriminative ability. Models were compared visually to determine if the refined TCellMap annotations provided superior predictive power over the baseline classifications.

### Cross-Validation analysis

To demonstrate whether predictive power was driven by a small subset of dominant features or the aggregate signature, we performed two additional experiments. First, I evaluated model stability across five discrete feature inclusion proportions (50%, 60%, 70%, 80%, and 90% of the granular sigNATURE features). For each proportion threshold, we randomly sampled the features and evaluated model performance using 5-fold cross-validation, repeating this entire process across 10 independent iterations per threshold.

Finally, to assess how predictive information was distributed across the latent structure of the data, we applied Principal Component Analysis (PCA) to the 27-dimension cellular proportion feature space and estimated the information gain contributed by each resulting principal component.

### Extension to the CD4+ T-Cell Compartment

To demonstrate the broader applicability of the analytical framework, we applied the identical pre-processing, sigNATURE label-transfer, and multivariate logistic regression pipelines described above to the CD4+ T-cell compartment isolated from the same Liu et al. clinical cohort. sigNATURE mapping and subsequent predictive modeling for the CD4+ cells were conducted using the exact same parameters and 5-fold cross-validation methodology utilized for the CD8+ compartment.

## DECLARATIONS

### Ethics approval and consent to participate

Not applicable.

### Consent for publication

Not applicable.

### Availability of data and materials

https://github.com/Sanikak24

### Authors’ contributions

Sanika Kamath implemented the method, conducted the experiments, and interpreted the results. Soyeon Kim discussed the results. Xueke Jin aided in setting up the code repository with Sanika. Hyun Jung Park and Jing Hong Wang conceived the project and interpreted the results; Hyun Jung Park designed the experiments and wrote the manuscript.

## Acknowledgements

This research was supported in part by the University of Pittsburgh Center for Research Computing, RRID:SCR_022735, through the resources provided. Specifically, this work used the HTC cluster, which is supported by NIH award number S10OD028483.

