## Supplementary figures and images for "sigNATURE maps cohort-specific T-cell states to reproducible programs of ICI response"

### S.Fig.1

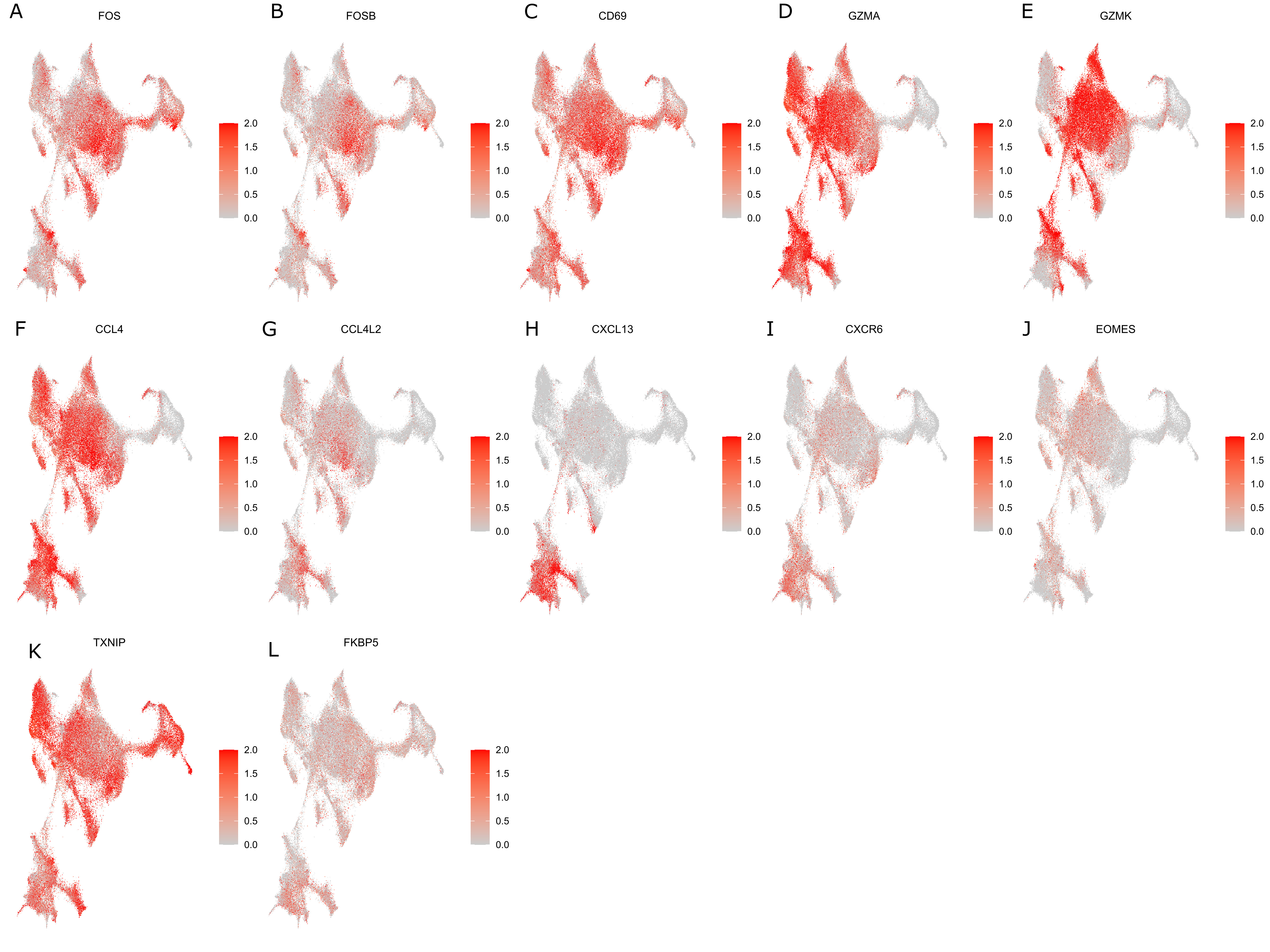

### S.Fig.2

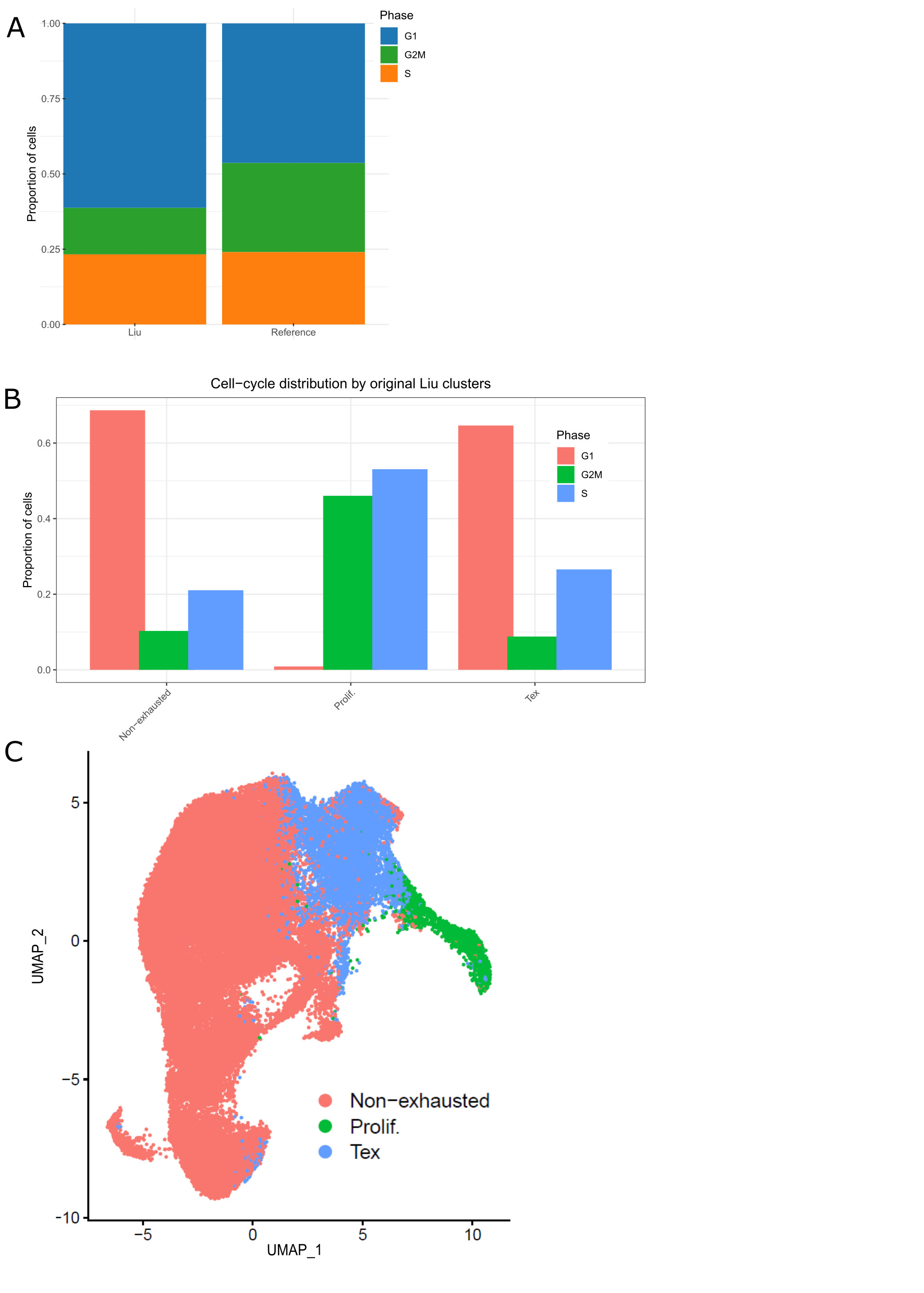

### S.Fig.3

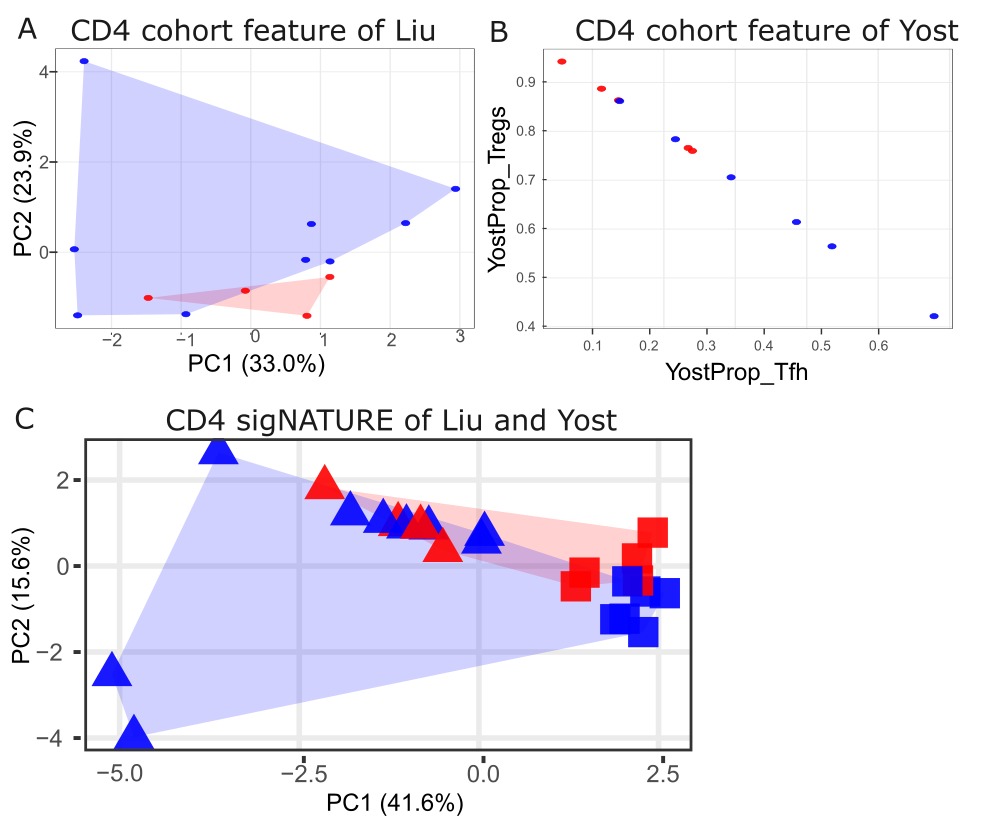
